# Full title:Age analysis and growth patterns of *Tenualosa ilisha* using otolith examination and length-weight relationships from different regions of Bangladesh

**DOI:** 10.1101/471664

**Authors:** Borhan Uddin Ahmed, A. K. Shakur Ahammad, Shahjahan, Biraj Kumar Datta, Fazla Rabbi, Mohammad Ashraful Alam, Abul Bashar, Yahia Mahmud

## Abstract

The assessment of age and growth patterns provides crucial information on numerous aspects of the population dynamics in fish, which can, in turn, be used to determine a sustainable fishing rate and inform effective resource management practices. However, studies on the age and growth pattern of *Tenualosa ilisha* (commonly referred to as hilsa) are plagued by a lack of essential information; addressing this data gap is the aim of the present study. Six diverse habitats across Bangladesh were chosen as sampling sites for hilsa collection. For age determinations, the lunar rings in the otolith of the hilsa fish that are periodically-deposited in accordance with the lunar cycle were used to reflect 14-day increments of time. The length and weight of each fish were recorded prior to otolith extraction. The resulting otoliths were polished to enable the visualization and quantification of their lunar rings with a high-magnification microscope. Except for the Kali River and Gaglajur Haor samples, the age of the fish correlated strongly with both their length and weight (*r* > 0.95; *p* < 0.05). Again, aside from those from the Kali River and Gaglajur Haor, all of the samples exhibited positive allometric growth patterns (*b* > 3) with the fish from the Tetulia River being the most positive (*b* = 3.48). The causes these variations are not yet clearly understood, however; the nutrient availability, environmental variation, and genetic-environmental interactions are likely contributors to the diversities displayed by the hilsa from different regions of Bangladesh.

## Introduction

*Tenualosa ilisha* (Fisher and Bianchi, 1984) the national fish of Bangladesh, is a member of Clupeidae family and a vital element of fish production in Bangladesh. It can also be referred to as ilish, hilsa, hilsa herring, or hilsa shad. It’s the largest single species of fish in Bangladesh and can be found in almost every river, estuary, and marine environment in the country [1], contributing 351 thousand metric tons (MT) [2] in total fish production annually. According to [3] hilsa represents approximately 1% of Bangladesh’s GDP with 500 – 600 million Tk. earning as foreign currency from the export of hilsa. It comprises 12% of the country’s total fish production [4]. Approximately 450,000 earn their livelihood by catching hilsa and 4 - 5 million people are indirectly involved in the industry through fisheries [5]. Hilsa can grow to up to 60 cm in length with weights of up to 4.2 kg [6]. The juvenile hilsa (measuring up to 25 cm) that return to the sea are known as “jatka” in Bangladesh. [7] studied the ecological distribution of hilsa and found that they are native to the foreshore areas, estuaries, brackish water lakes, and freshwater rivers of the western division of the Indo-Pacific region. Its marine distribution extends from the Persian Gulf near Iran and Iraq to the Arabian Sea and the Bay of Bengal on India’s west coast. It has also been reported to inhabit the coastal waters of Sri Lanka and Cochin, China (Laos). Currently, the upstream population has been severely depleted and the fish are mainly concentrated in the downstream rivers, estuaries, coastal areas, and Bay of Bengal [8]. There are many factors involved in the decline of the hilsa population in Bangladesh including the indiscriminate exploitation of the juvenile (Jatka) and smaller-sized hilsa for consumption, reduced water influx due to the Farakka barrage, increased river pollution and silt deposition, destruction of their migration routes, spawning, feeding and nursing grounds by human intervention. Hence, there is growing concern among marine biologists surrounding the conservation and sustainable maintenance of this species, which may soon require imposing different management strategies, regulations, and interventions.

When evaluating the stock of any fish species, many different population parameters (e.g. age, growth rate, sex, reproduction, etc.) must be taken into consideration. Information on age and growth is extremely important to almost every aspect of fisheries [9]. However, recently an alarming phenomenon has been observed in Bangladesh, namely, the availability of small-sized hilsa with mature gonads. It remains unknown whether these are adult fish or whether environmental factors (e.g. climate, hydrological, and ecological changes) have induced premature gonad maturation in the juveniles [10]. Also, sexual maturity can be reached at different sizes and ages due to spawning migration as the fish takes a lot of food during their migration [11-12]. Differences in mortality, growth rates, and long life span can also cause sexual maturity in this fish [13]. Therefore, potential explanations for the early gonadal development observed in hilsa involve genetic factors arising from inbreeding depression, densely populated areas, food competition, and incomplete migration. Some small-sized hilsa may remain in the river rather than return to sea and, thus, reach sexual maturity in this environment. Determining the age of these fish is crucial for testing this hypothesis and therefore, is the focus of the present study. For age determination, different researchers have used different methods that rely on the analysis of the scales, otoliths, fin spines, etc. But because these are all external structures that may be shed and regenerated anew over the course of the fish’s life cycle, they do not reliably represent the age of the fish. In contrast, the otolith is an internal organ that never detaches from the body of the fish. This characteristic renders the otolith ring particularly well suited for lifespan studies and was therefore chosen for the present study. Otoliths are paired calcified structures used for balance and/or hearing in teleost fish. The otolith has long been known as a timekeeper in fish. Technological developments over the past few decades have facilitated the analysis of otoliths and the information gained from these experiments has greatly expanded our understanding of these fish [14-17]. Otoliths are naturally-occurring data loggers that use microstructures to record different temporal events related to the growth and environment experienced by the fish [18]. Moreover, the otolith also encodes crucial information about fish’s age, growth rate, movement patterns, and habitat interactions that can be extrapolated to the population level in terms of the ecology, demography, and life history of the species [19]. No other biological structure even compares to the otolith in terms of ability to relay information on the age and growth patterns of the fish [20]. However, the otolith in *T. ilisha* does not contain yearly rings, also referred to as annuli, which makes using this structure to determine the age more challenging in this species. However, lunar rings are present in the otolith of the hilsa and these also accurately and reliably reflect the age of the fish [21]. In the present study, lunar rings were used to age hilsa collected from different regions of Bangladesh. This approach has not yet been described in the literature and, therefore, the present study provides novel insight into the relationship between the age and size of Bangladesh’s most important species of fish.

## Materials and Methods

### Sample collection and otolith extraction

Fresh hilsa were collected randomly from six different locations in Bangladesh: Meghna River Estuary Chandpur; Bay of Bengal, Cox’s Bazar; Kali River, Kishoreganj; Tetulia River, Barisal; Padma River, Mawa, Munshiganj; and Gaglajur Haor, Mohanganj, Netrokona (Fig 1). Approximately 30 samples were collected from each area. Otolith extraction was performed immediately upon collection of the fresh fish (S1 Fig). Prior to this dissection, the total length and weight of each fish were recorded in cm and g, respectively. The isolated otoliths were washed, air-dried, and stored in a 1-ml Eppendorf tube in 100% ethanol at room temperature.

**Fig 1.**
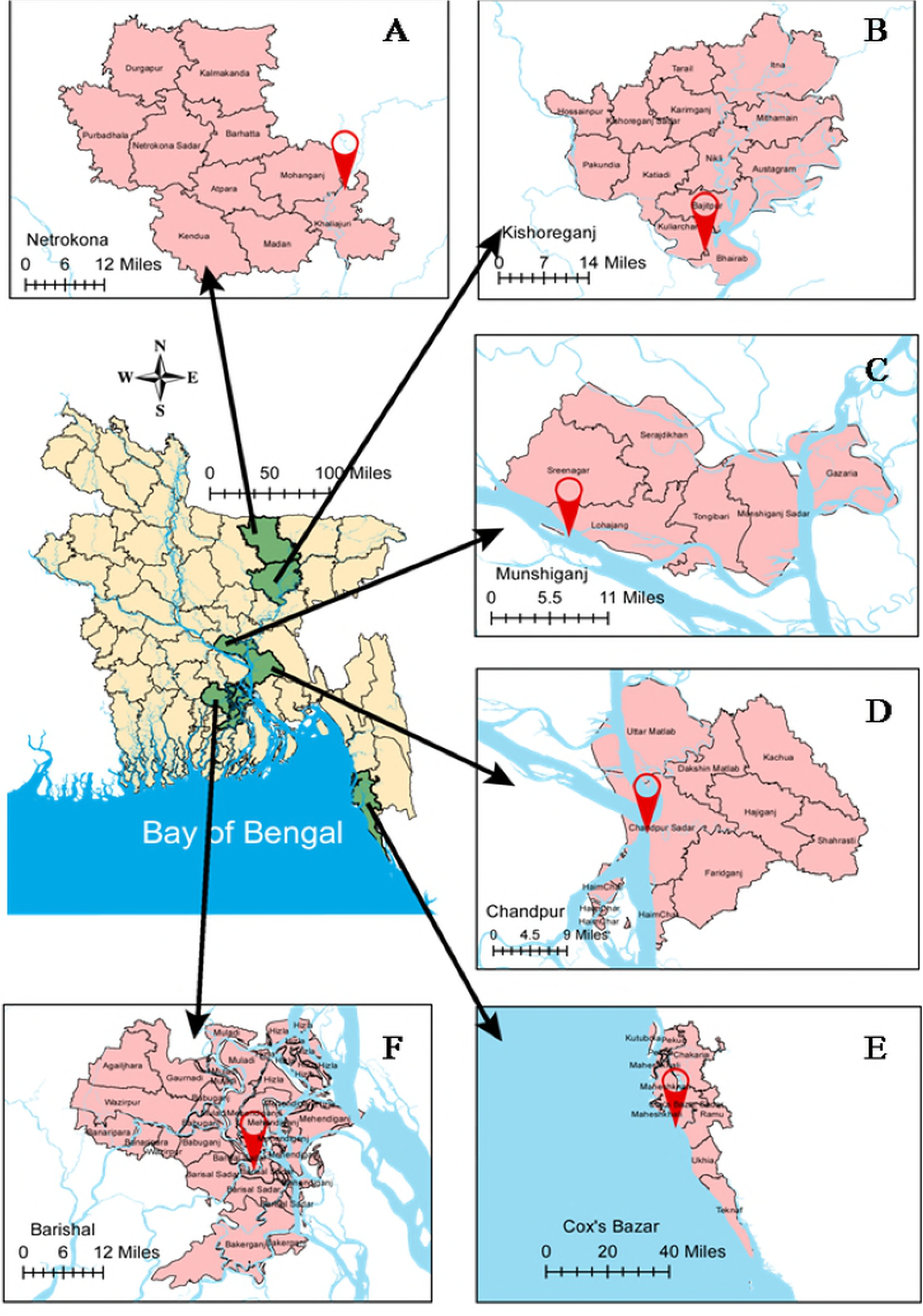
Hilsa sampling sites from different region of Bangladesh. (A) Gaglajur haor, Mohanganj, Netrokona; (B) Kali river, Kishoreganj; (C) Padma river, Mawa, Munshiganj; (D) Meghna river estuary, Chandpur; (E) Bay of Bengal, Cox’s Bazar; (F) Tetulia river, Barishal

### Otolith processing and observation

To visualize the rings in the otolith, they were ground and polished using a combination of the acid contact method [22] and the wet-dry sandpaper method [21]. For the acid contact method, the otoliths were first submerged into distilled water for one minute. Their convex exterior was then ground by hand against a fine carborundum stone the surface of which was coated in dilute hydrochloric acid. The use of too much acid was avoided to not allow the edge of the otolith to contact the acid. During this grinding procedure, the otolith was sporadically examined under a microscope to check for the appearance of the rings. When the rings became clearly visible, the otolith was again submerged into distilled water for 1 to 2 minutes to remove the acid, followed by 70% ethanol, and then absolute ethanol for the following 1 to 2 minutes. The wet-dry sandpaper method utilized pieces of sandpaper with grit sizes that ranged from 600 to 1000. Using a spherical motion with the sandpaper ensured that every side of the otolith was polished evenly. This process was carried out until the rings became observable using a microscope and then otolith was washed in distilled water and allowed to air dry. The rings were counted under a light microscope at 100x magnification using immersion oil. Any otoliths without clearly-visible rings were regarded as too poorly-polished to be included in future analyses.

### Age calculation

Both the daily and lunar rings were visible under 100x magnification. According to [21] lunar rings consist of 11 serially-deposited, comparatively light, narrow rings, followed by three deeper, wider rings These three rings are likely formed during the spring tide, over the course of the three most active feeding and migration days for the hilsa [23]. The deeper, wider rings appear as one thick, dark band, referred to as a lunar ring, and are indicative of a 14-day increment of time. The 14-day periodical development of the lunar rings occurs in accordance with the lunar cycle. The validity and reliability of using these chronological markings to calculate the age of a fish were verified by quantifying the rings in fish raised in captivity for known periods of time [24] and by marginal increment analysis [25]. In the present study, less than 10% unreadable area of otolith (estimated by Sigma Plot.V.13) were accepted for ring counting. The 11 daily rings were not clearly visible in a few but not all of the otoliths. Therefore, only the lunar rings were included in our age calculation, which was generated using the following formula (Equation 1):

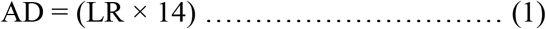

Where AD is the age of the hilsa in days and LR is the total number of lunar rings counter in the otolith.

### Length calculation

According to [26], regression relationships effectively associate length with age. In the present study, a linear regression model was established between the fish’s length and its age (determined from Equation 1) using the following formula (Equation 2):

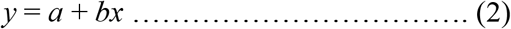

Where *y* is the total length (cm); *x* is the age (months); *a* is the intercept of the linear regression; and *b* is the slope.

### Length-weight relationship calculation

The relationships between length and weight were calculated for all of the samples using the cube law given by [27] (Equation 3)

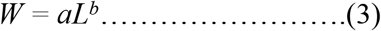

Where *W* is the body weight (g); *L* is the total length (cm); *a* is the regression intercept, and *b* is the regression slope. of the graph. When *b* is equal to 3, the growth of the fish is considered to be isometric [28-29]

### Statistical analysis

The data obtained from the length, weight, and lunar rings of the hilsa were analyzed using SPSS v. 21 (IBM SPSS for windows, Armonk, New York, USA) and Microsoft Excel program v. 2016. All data were subjected to an analysis of variance (ANOVA) followed by a comparison of the means using Duncan’s multiple range test at a 95% confidence level.

## Results

### Otolith structure

The otoliths of *T. ilisha* were laterally compressed and whitish in color (S2 Fig). One surface was convex and the other concave. The point at its center was encircled by the rings that served as the metric for calculating the age of the fish. The anterior margin branched into the rostrum and anti-rostrum, separated by a wide, deep excisural notch, and the rostrum was smaller than the anti-rostrum (S2 Fig). The posterior portion of the otolith had a rough edge where anterior part found to be smooth.

### Age determination

The age of each hilsa was calculated by counting the lunar rings found in their otolith (Fig 2). Approximately two out of every three otoliths were sufficiently polished and quantified. Data obtained from these calculations indicated that the age of these fish ranged from one to over five years old. The average total lengths were as 29.50 ± 1.76 cm, 34.28 ± 3.34 cm, 40.66 ± 1.96 cm, 44.6 ± 2.30 cm, and 45.55 ± 0.35 cm for the fish aged between 1-2 years, 2-3 years, 3-4 years, 4-5, and 5-6 years, respectively (Table 1). The relationship between length and age was linearly proportional in all of the fish collected from the six sampling sites; this indicates that the length of the hilsa increases according to its age independent of its habitat (Fig 3). Similarly, the relationship between the hilsa’s weight and its age was also preserved across all of the samples (Fig 4). The results obtained from counting the lunar rings in the otolith were used to establish linear regression equations describing the length-age and the weight-age relationship of the hilsa from each sampling site (*y* = *a* + *bx*), where the slope value *b* describes the increase in length and weight associated with each month (Figs. 3 and 4). In all of the samples, both the length-age and weight-age correlations were statistically significant (*r* > 0.95; *p* < 0.05) except for those collected from the Kali River and Gaglajur Haor (*r* < 0.95 and *p* > 0.05) (Figs. 3 and 4).

**Fig 2.**
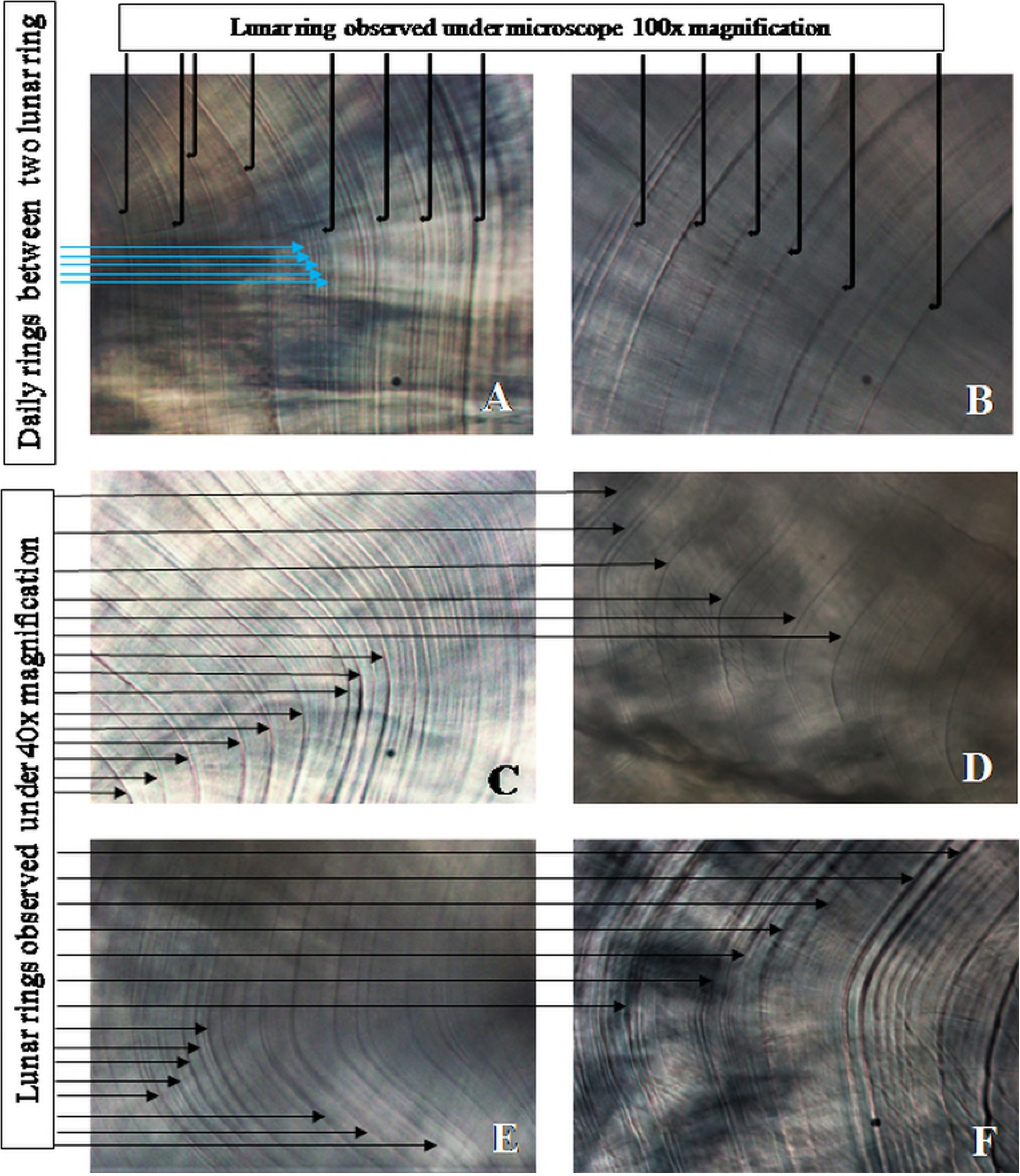
Lunar rings from the sagittal otolith of *Tenualosa ilisha*. (A) Daily and (B) lunar rings visualized using a compound microscope (Olympus CX-41, Shinjuku, Tokyo, Japan.) at 100x magnification; (C) – (F) Lunar rings observed at 40x magnification.

**Table 1:** The average length, weight, number of rings and the calculated age grouped according to size

| Size group | N | Average length (cm) | Average weight (g) | Average lunar ring (n) | Calculated age |  |
| --- | --- | --- | --- | --- | --- | --- |
|  |  |  |  |  | Month | Year |
| Small | 66 | 29.50± 1.76 | 265.52±43.45 | 46±7.40 | 21 | > 1 |
| Medium | 59 | 34.28±3.34 | 461.27±110.94 | 61±7.49 | 28 | > 2 |
| Large | 42 | 40.66±1.96 | 742.75±78.36 | 87±12.59 | 40 | > 3 |
| Intermediate-large | 9 | 44.6±2.30 | 920±81.66 | 109±8.44 | 50 | > 4 |
| Extra-large | 4 | 45.55±0.35 | 1372.5±77.07 | 142±1.41 | 65 | > 5 |

**Fig 3.**
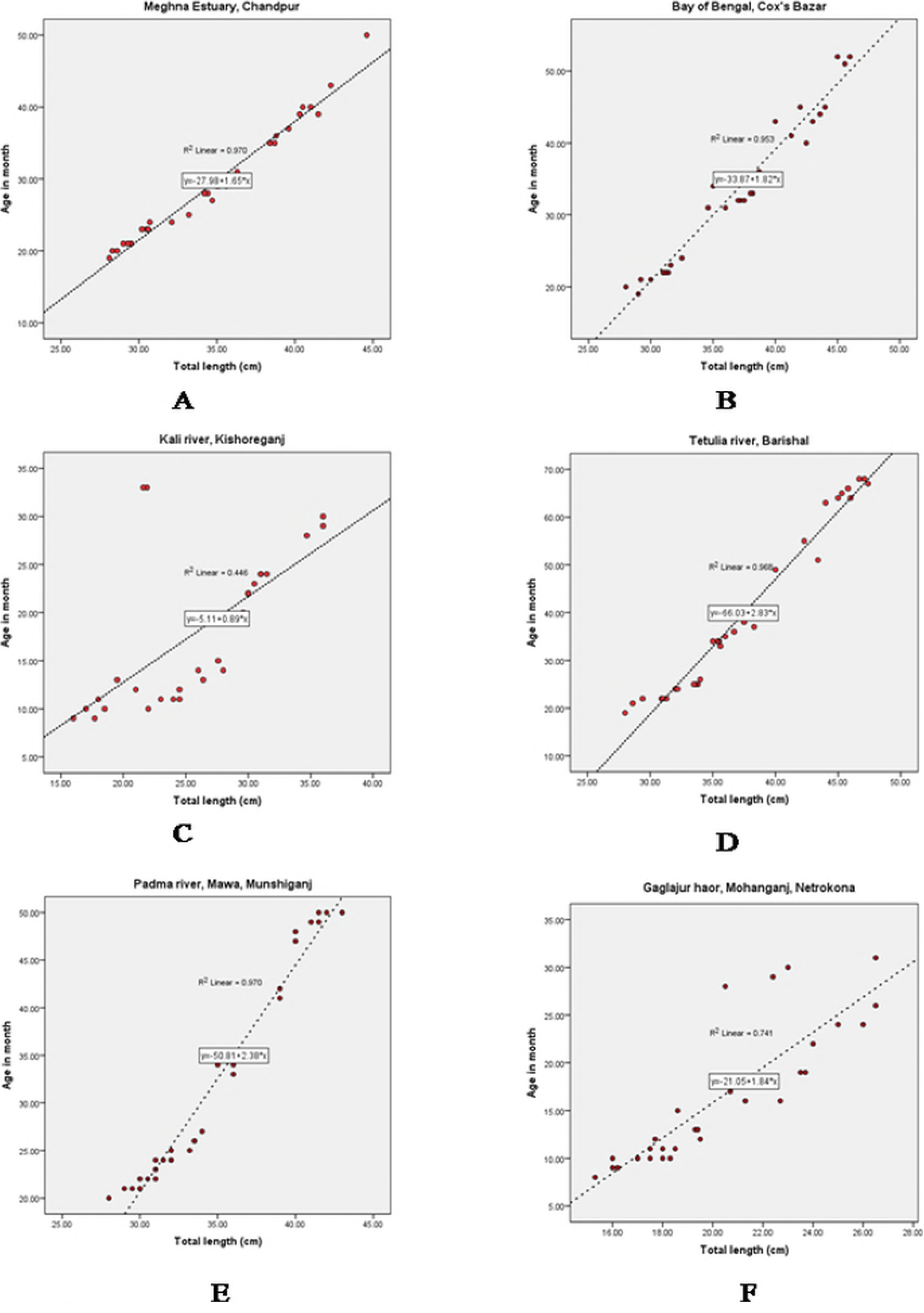
Relationships between the total length (cm) and age (month) of *Tenualosa ilisha* from six different areas of Bangladesh. (A) Meghna River Estuary, Chandpur; (B) Bay of Bengal, Cox’s Bazar; (C) Kali River, Kishoreganj; (D) Tetulia River, Barishal; (E) Padma River, Mawa, Munshiganj; (F) Gaglajur Haor, Mohongonj, Netrokona. All values were generated from otolith readings and combined for both sexes.

**Fig 4.**
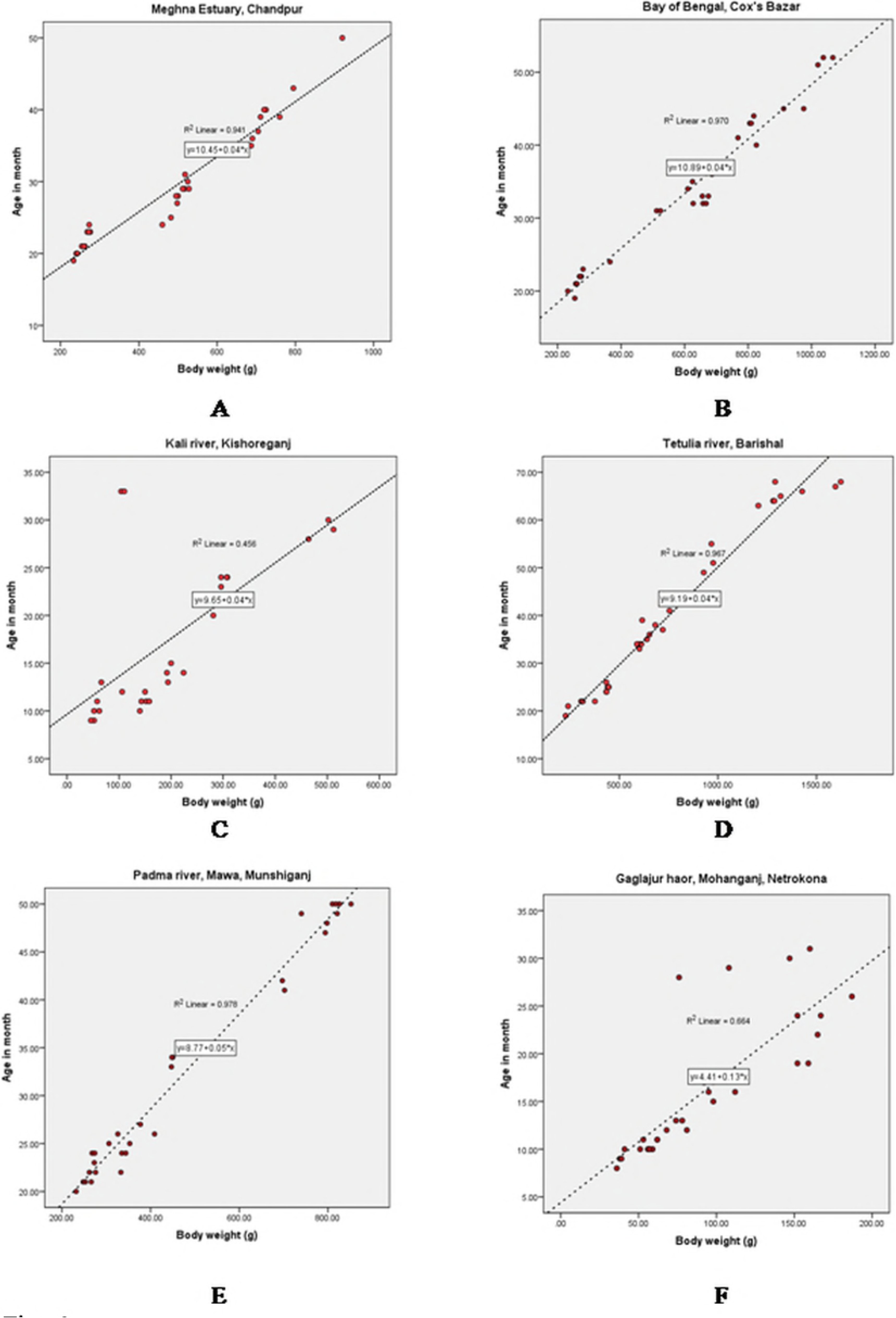
Relationships between the body weight (g) and age (months) of *Tenualosa ilisha* from six different areas of Bangladesh. (A) Meghna River Estuary, Chandpur; (B) Bay of Bengal, Cox’s Bazar; (C) Kali River, Kishoreganj; (D) Tetulia River, Barishal; (E) Padma River, Mawa, Munshiganj; (F) Gaglajur Haor, Mohongonj, Netrokona. All values were generated from otolith readings and combined for both sexes.

### Length-weight relationship

Calculating the length-weight relationship of each sample using Equation 3 revealed that the highest slope value (*b*) was recorded from the Tetulia River (*b* = 3.5), followed by the Padma River (*b* = 3.4), Bay of Bengal (*b* = 3.3), Meghna River Estuary (*b* = 3.2), Kali River (*b* = 3.0), and Gaglajur Haor (*b* = 3.0) (Fig 5). The results describe positive allometric growth behavior in all of the samples except for those from the Kali River and Gaglajur Haor, which exhibited isometric and negative allometric growth patterns, respectively. The results obtained from the length-weight relationship analysis and their associated descriptive statistics are presented in Table 2 for all of the hilsa according to sampling site.

**Fig 5.**
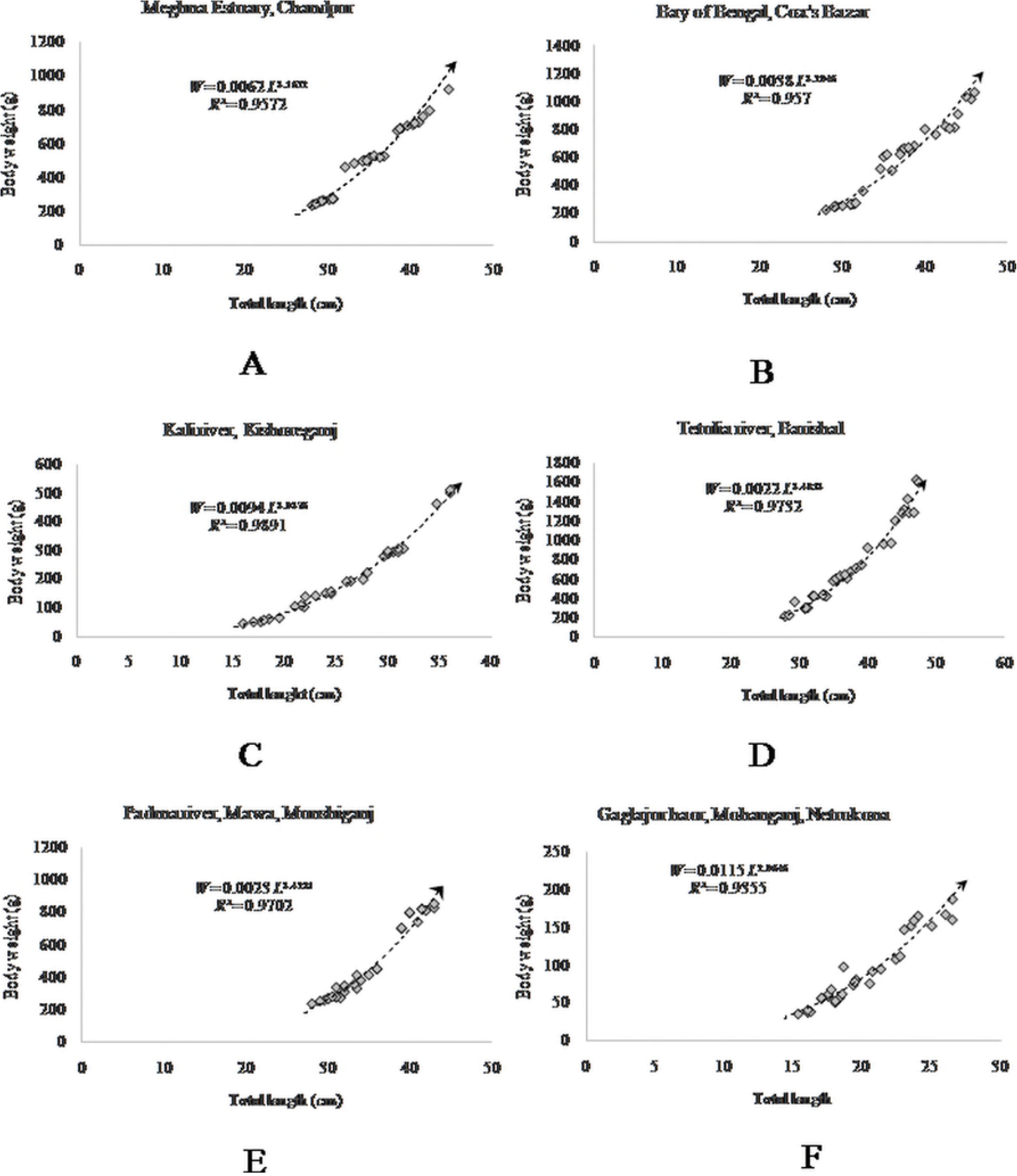
Length-weight relationship in *Tenualosa ilisha* from six different areas of Bangladesh. (A) Meghna River Estuary, Chandpur; (B) Bay of Bengal, Cox’s Bazar; (C) Kali River, Kishoreganj; (D) Tetulia River, Barishal; (E) Padma River, Mawa, Munshiganj; (F) Gaglajur Haor, Mohongonj, Netrokona. Values represent those from both sexes combined.

**Table 2.** Length-weight relationship parameters and associated descriptive statistics.

| Collection site | Weight<br>(mean±SD) | Length<br>(mean±SD) | Determination<br>coefficient ($r^2$) | Slope<br>value<br>($b$) | Growth<br>pattern |
| --- | --- | --- | --- | --- | --- |
| Meghna River<br>Estuary | 500.17±204.26 | 34.93±4.82 | 0.95** | 3.1622 | (+)<br>Allometric |
| Bay of Bengal | 582.33±252.04 | 36.85±5.54 | 0.93** | 3.1717 | (+)<br>Allometric |
| Kali River | 211.46±131.49 | 25.92±5.77 | 0.98* | 3.0376 | Isometric |
| Tetulia River | 750.78±411.56 | 37.49±5.97 | 0.97** | 3.4813 | (+)<br>Allometric |
| Padma River | 476.60±230.95 | 34.95±4.78 | 0.97** | 3.4231 | (+)<br>Allometric |
| Gaglajur Haor | 92.73±46.52 | 20.19±3.38 | 0.93* | 2.9646 | (-)<br>Allometric |

\*\* $P < 0.01$ ; \* $P < 0.05$

## Discussion

The external anatomy of the otolith from *T. ilisha* determined in this study are similar to those described by [1] and comparable to other species of clupeid fish characterized by [30]. The otoliths investigated here were slightly convex or oval in shape that made polishing them challenging. When the top portion was properly polished, the edges were not; yet, any further polishing would cause the rings in the top portion to disappear from over-polishing just when the rings around the edges became visible. Hence, the otolith from *T. ilisha* proved to be very difficult to polish. Annulus formation in fish is generally thought to coincide with the spawning season [31]. Furthermore, the impetus for annulus formation and its timing have been associated with gonadal maturation and may be partially dependent on nutrient availability, starvation, environmental variation, and stress [32]. In fish, starvation generally occurs at extremely low temperatures. In Bangladeshi waters, temperature drops are not detrimental enough to completely cease feeding. Therefore, a partial break in growth is very rare for hilsa, which could explain the absence of annuli in the otolith of this species of fish. However, both migration and the formation of lunar rings occur according to the lunar cycle and hilsa are highly migratory.

The calculated ages and corresponding age groups determined in the present study are in agreement with those described by [33] who estimated the age of hilsa using length frequency analysis to be between 2-6 years using a population in which 90% were between 25-75 cm in length. The length and weight at age findings obtained in the present study (except for those collected from the Kali River and Gaglajur Haor) support the results presented in [34] from hilsa from Indian water. In addition, [31] reported a similar relationship with age using otolith ring analysis of fish from the Padma and Meghna rivers. [35] also conducted a study on the daily rings in the otoliths from hilsa and reported similar results in terms of length at age as were generated here. However, the length at age values from all of the sites in the present study were slightly lower than those reported by [1]. The variations between age and length or weight correlations from the Kali River and Gaglajur Haor samples may be due to their separate environments because this two sampling site is in upstream portion of the country which may cause difference of availability of nutrients at these two sites from other four. The variations could also be reflective of genetic issues, for example, inbreeding, genetic depression, or changes to the expression levels of genes related to growth. The size differences observed between similarly-aged hilsa could result from habitat-associated gene expression changes in addition to climate change, food availability, or a combination of these factors. Identifying the underlying cause of this phenomenon unequivocally requires studies that include a high-resolution examination of the genes related to gonad development (e.g. *Dmrt, Foxl2*, etc.).

In terms of the length-weight relationship, a study conducted by [36] in the Tetulia River also found positive allometric growth patterns for both male (*b* = 3.02) and female (*b* = 3.08) hilsa and strong positive correlations between their length and weight (male; *r^2^* = 0.969. female; *r^2^* = 0.968), which supports the results obtained in the present study (again, except for the Kali River an Gaglajur Haor samples). In contrast, [37] observed growth patterns in fish from the Meghna Estuary of Chandpur that they characterized as isometric after analyzing the length-weight relationship; these findings disagree with the positive allometric growth found in the present study. This variation is likely explained by differences in the study periods and the resulting changes to food availability, the climate, and, potentially, gene-environmen interactions.

## Conclusions

Evidence-based assessment of fish age is necessary for stock assessments and to develop appropriate management strategies and effective conservation plans. Spawning, feeding, migration, and other physiological activities of the fish are greatly influenced by their age. The early maturation of the hilsa in Bangladesh is also associated with their age. Data on the age of the hilsa provides a basis for calculating their growth rate, mortality, and productivity of this species. Therefore, age is an important parameter that can inform choices related to the sustainable and effective management of the hilsa population. Taken together, the results provided by the present study will help other researchers to conduct further research in this area and enable policy makers to develop appropriate strategies that ensure the hilsa population maintains its position in the economy, culture, and cuisine of Bangladesh.

## Acknowledgments

The authors would like to acknowledge the Bangladesh Fisheries Research Institute (BFRI) for providing funds for this research. The authors also would like to thank Dr. Yahia Mahmud, Director General, BFRI, for access to laboratory facilities and a research vessel as well as M.V. Rupali ilish for help with sample collection.

## Conflict of interest

The authors have no conflicts of interest. The funding agency had no role in the design of the study; in the collection, analyses, or interpretation of data; in the writing of the manuscript; or in the decision to publish the results.

## Supporting information

**S1 Fig.**
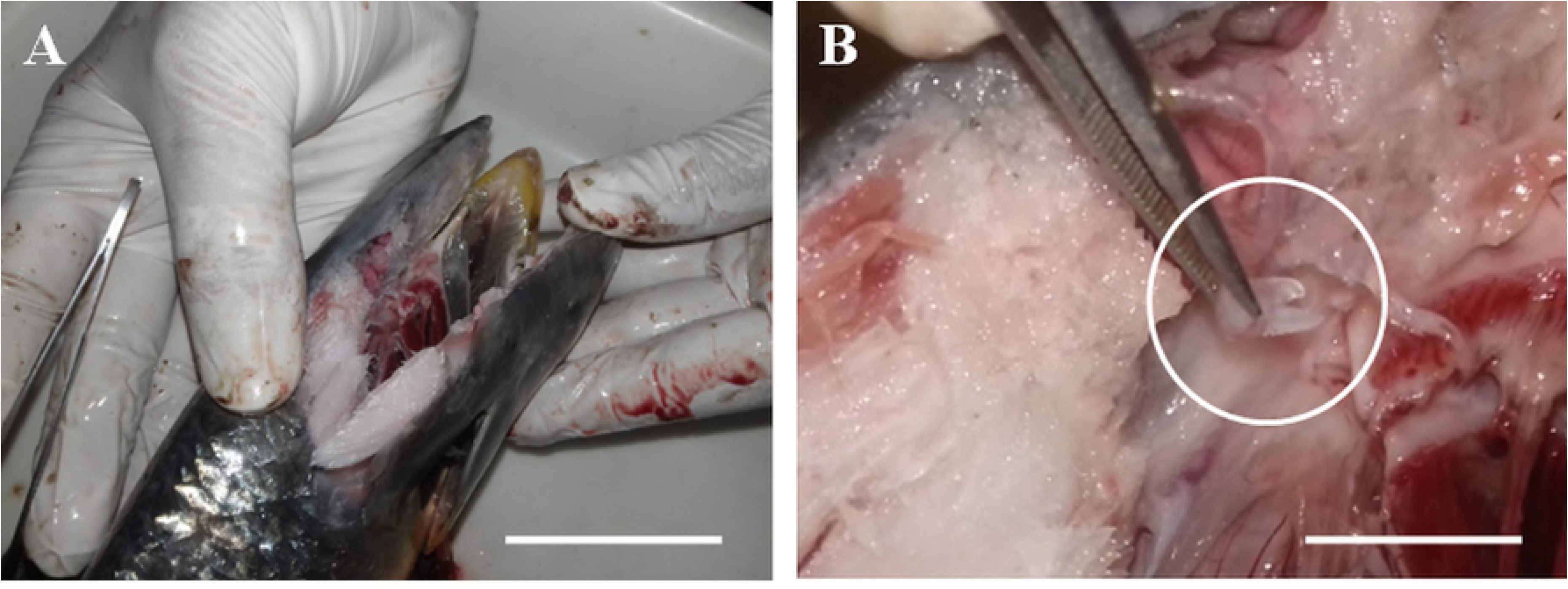
Head dissection and otolith collection. (A) Dissection of the hilsa’s head using a knife and scalpel. (B) Collection of the sagittal otolith from the dissected head.

**S2 Fig.**
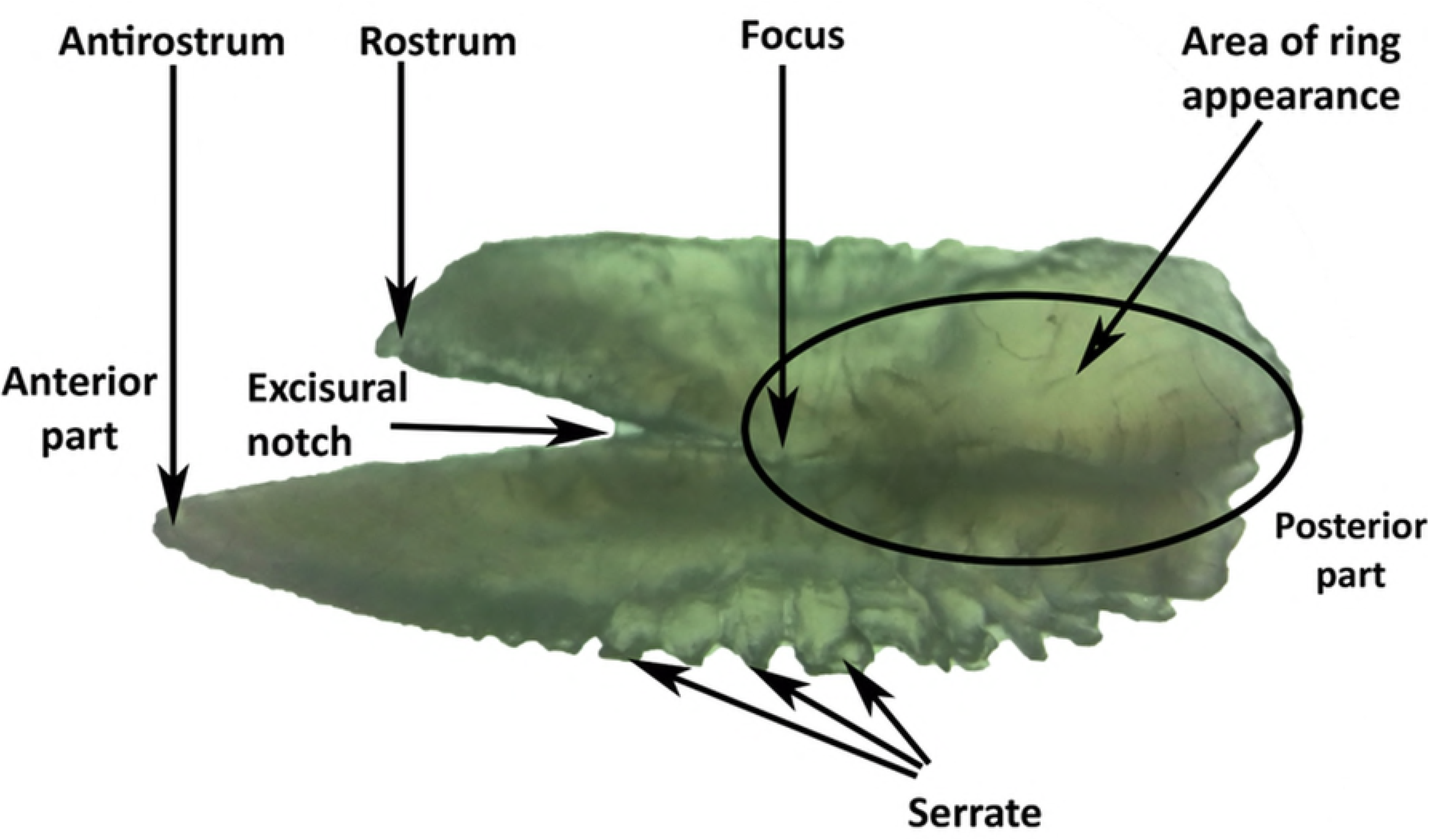
Otolith structure in hilsa. Different black arrow indicates different parts of the segital otilith of *T. ilisha*.

**Author contributions**
Conceptualization, A.K.S.A.; supervision, A.K.S.A.; methodology, M.B.U.A.; manuscript editing, M.S.; sample collection, B.K.D; data arrangement, M.F.R; investigation, M.A.A.; funding acquisition, A.B.; writing - original draft preparation, M.B.U.A.; writing - review and editing, M.A.A.; project administration, Y.M.

## References

1. Ahmed MS, Sharif ASM, Latifa GA. Age, Growth and Mortality of Hilsa Shad, *Tenualosa ilisha* in the River Meghna, Bangladesh. Asian Journal of Biological Sciences. 2008; 1:69–76.

2. DoF (Department of Fisheries). National fish week, compendium (In Bengali). Department of Fisheries, Ministry of Fisheries and Livestock, Dhaka, Bangladesh. 2014.

3. Bala BK, Arshad FM, Alias EF, Sidique SF, Noh KM, Rowshon MK, et al. Sustainable exploitation of hilsa fish (*Tenualosa ilisha*) population in Bangladesh: Modeling and policy implications. Ecological modelling. 2014; 283:19–30.

4. Rahman MA, Alam MA, Hasan SJ, Jaher M. Hilsa fishery management in Bangladesh. Hilsa: Status of fishery and potential for aquaculture. 2012; pp 40–60.

5. DoF (Department of Fisheries). National fish week, compendium (In Bengali). Department of Fisheries, Ministry of Fisheries and Livestock, Dhaka, Bangladesh. 2015.

6. Bhaumik U, Mukhopadhyay MK, Shrivastava NP, Sharma AP. The largest recorded Hilsa (*Tenualosa ilisha*) in India from Tapti estuary, Gujarat. Fishing Chimes. 2012; 31(12):57–58.

7. Pillay SR, Rosa Jr H. Synopsis of biological data on hilsa, *Hilsa ilisha* (Hamilton, 1882). FAO Fisheries Biology Synopsis. 1963; 25 (1):1–6.

8. Haldar GC, Rahman MA. Ecology of hilsa, *Tenualosa ilisha* (Fisher and Bianchi, 1984)). In Proceedings of BFRI/ACIAR/CSIRO Workshop on Hilsa Fisheries Research in Bangladesh, held on 3-4 March, 1998 at Bangladesh Agricultural Research Council, Dhaka, Bangladesh. 1998; pp. 3–4.

9. Jobling M. Environmental factors and rates of development and growth. Handbook of Fish Biology and Fisheries. 2002; 1:97–122.

10. Almukhtar MA, Jasim W, Mutlak F. Reproductive Biology of Hilsa Shad *Tenualosa ilisha* (Teleostei: Clupeidae) During Spawning Migration in the Shatt Al Arab River and Southern Al Hammar Marsh, Basra, Iraq. Journal of Fisheries and Aquatic Science. 2016; 11(1):43.

11. Mohamed ARM, Qasim AM. Stock assessment and management of hilsa shad (*Tenualosa ilisha*) in Iraqi marine waters, northwest Arabian Gulf. International. Journal of Fisheries and Aquatic Study. 2014; 1(5):1–7.

12. Hussain SA, Al–Mukhtar MA, Al–Daham NK. Preliminary investigation on fisheries and some biological aspects of sbour, *Hilsa ilisha* from Shatt al–Arab River, Iraq. Basrah Journal of Agricultural Science. 1991; 4(1 and 2): 141–151.

13. Zhang J, Takita T, Zhang C. Reproductive biology of *Ilisha elongata* (Teleostei: Pristigasteridae) in Ariake Sound, Japan: Implications for estuarine fish conservation in Asia. Estuarine, Coastal and Shelf Science. 2009; 81(1): 105–113.

14. Grant A. Age determination and growth in fish and other aquatic animals. Australian Journal of Marine and Freshwater Research. 1992; 43: 879–1330.

15. Campana SE, Secor DH, Dean JM. Recent developments in fish otolith research. University of South Carolina Press. 1995; pp. 89–99.

16. Panfili J, De Pontual H, Troadec H, Wrigh PJ. Manual of fish sclerochronology. (IFREMER-IRD) 1^st^ Ed^n^., Brest, France. 2002.

17. Campana SE, Thorrold SR. Otoliths, increments, and elements: keys to a comprehensive understanding of fish populations? Canadian Journal of Fisheries and Aquatic Sciences. 2001; 58(1): 30–38.

18. Kalish JM. Otolith microchemistry: validation of the effects of physiology, age and environment on otolith composition. Journal of Experimental Marine Biology and Ecology. 1989; 132(3): 151–178.

19. Campana SE. Chemistry and composition of fish otoliths: pathways, mechanisms and applications. Marine Ecology Progress Series. 1999; 188: 263–297.

20. Begg GA, Campana SE, Fowler AJ. (Eds.). Fish Otolith Research and Applications: Proceedings of the Third International Symposium on Fish Otolith Research and Application, Townsville, Queensland, Australia, 11-16 July 2004. CSIRO.

21. Rahman MJ, Cowx IG. Lunar periodicity in growth increment formation in otoliths of hilsa shad (*Tenualosa ilisha*, Clupeidae) in Bangladesh waters. Fisheries Research. 2006; 81(2-3):342–344.

22. Sinha VRP, Jones JW. On the age and growth of the freshwater eel (*Anguilla anguilla*). Journal of Zoology. 1967;153(1): 99–117.

23. Rahman MJ. Population biology and management of hilsa shad (*Tenualosa ilisha*) in Bangladesh (Doctoral Dissertation) University of Hull, England. 2001; p. 253.

24. Hayashi A, Kawaguchi K, Watanabe H, Ishida M. Daily growth increment formation and its lunar periodicity in otoliths of the myctophid fish *Myctophum asperum* (Pisces: Myctophidae). Fisheries Science. 2001; 67(5): 811–817.

25. Gartner JV. Life histories of three species of lantern fishes (Pisces: Myctophidae) from the eastern Gulf of Mexico. Marine Biology. 1991; 111(1): 11–20.

26. Campana SE, Jones CM. Analysis of otolith microstructure data. Otolith microstructure examination and analysis. Edited by DK Stevenson and SE Campana. Canadian Special Publication of Fisheries and Aquatic Science. 1992; 117: 73–100.

27. Le Cren ED. The length-weight relationship and seasonal cycle in gonad weight and condition in the perch (*Perca fluviatilis*). The Journal of Animal Ecology. 1951; 201–219.

28. Edwards AL. An introduction to linear regression and correlation. W.H. Freeman and Company, USA. 1976; p. 213.

29. Draper NR, Smith H. Applied Regression Analysis. Wiley Series in Probability and Mathematical Statistics, John Wiley and Sons. New York, USA. 1981; p. 709.

30. Smale MJ, Watson G, Hecht T. Otolith Atlas of Southern African Marine Fishes. Ichthyological Monographs, South Africa. 1995; 1:1–253.

31. Methot Jr RD. Seasonal variation in survival of larval northern anchovy, *Engraulis mordax*, estimated from the age distribution of juveniles. Fishery Bulletin. U.S.A. 1983; 81: 741–750.

32. Pannella G. Growth patterns in fish sagittae. In, Skeletal growth of aquatic organirms: biological records of environmental changes, edited by D.C. Rhoads and R. A. Lutz, Plenum Press, New York. 1980; p. 519–560.

33. Hossain MAR, Das I, Genevier L, Hazra S, Rahman M, Barange M. et al. Biology and fisheries of Hilsa shad in Bay of Bengal. Science of the Total Environment. 2018; 651(2):1720–1734.

34. De DK, Datta NC. Age, growth, length-weight relationship and relative condition in hilsa, *Tenualosa ilisha* (Fisher and Bianchi, 1984) from the Hooghly estuarine system. Indian Journal of Fisheries. 1990; 37(2): 199–200.

35. Blaber SJM, Milton DA, Chenery SR, Fry G. New insights into the life history of *Tenualosa ilisha* and fishery implications. American Fisheries Society Symposium. 2003; 35: 223–240.

36. Karim R, Roy KC, Roy PR, Ahmed ZF. Age and growth of hilsa shad, *Tenualosa ilisha* (Hamilton, 1822) of the river Tentulia in Bangladesh. Journal of Fisheries. 2015; 3(1): 227–232.

37. Flura MZ, Rahman BS, Rahman MA, Alam MA, Pramanik MH. Length-weight relationship and GSI of hilsa, *Tenualosa ilisha* (Fisher and Bianchi, 1984) fishes in Meghna river, Bangladesh. International Journal of Natural and Social Science. 2015; 2(3):82–88.

